# Epidemic patterns of emerging variants with dynamical social distancing

**DOI:** 10.1101/2023.02.03.526970

**Authors:** Golsa Sayyar, Gergely Röst

## Abstract

Motivated by the emergence of new variants during the COVID-19 pandemic, we consider an epidemiological model of disease transmission dynamics, where novel strains appear by mutations of the virus. In the considered scenarios, disease prevalence in the population is modulated by social distancing. We study the various patterns that are generated under different assumptions of cross-immunity. If recovery from a given strain provides immunity against all previous strains, but not against more novel strains, then we observe a very regular sequential pattern of strain replacement where newer strains predominate over older strains. However, if protection upon recovery holds only against that particular strain and none of the others, we find much more complicated dynamics with potential recurrence of earlier strains, and co-circulation of various strains. We compare the observed patterns with genomic analysis we have seen during the COVID-19 pandemic.

## 1 Introduction

COVID-19 is a highly contagious viral disease caused by the severe acute respiratory syndrome coronavirus 2 (SARS-CoV-2). The disease quickly spread worldwide, resulting in the COVID-19 pandemic that has been declared the most significant global health crisis since the influenza pandemic of 1918 [1]. Since it was initially detected in late December 2019 in Wuhan, Hubei Province, China; as of October 2022, more than 600 million confirmed cases, including over 6.5 million deaths, have been reported on the World Health Organization (WHO) coronavirus (COVID-19) dashboard.

Mathematical models have been commonly applied throughout the pandemic to inform decision-makers and help the pandemic response. In the early phase, the potential for the global spread was the main concern [2, 3]. Control measures, including social distancing measures that reflect a strong effort to suppress or at least slow down the spread of the virus, began in mid-March 2022 in most European countries. When countries introduced non-pharmaceutical measures, mathematical models were used to evaluate the various disease control strategies [4, 5]. Compartmental models describing the transmission dynamics of the infection, have been extended by considering social distancing measures as manipulable control inputs to devise intervention strategies with optimal timing and intensity [6]. Mathematical models were also useful to give insights into the potential and optimization of testing strategies [7], including group testing [8]. The impact of vaccination and the waning and boosting of immunity have also been evaluated by models [9, 10]

With the emergence of new strains of the virus, researchers considered the spread of multiple lineages [11], the potential for new waves [12], and the adaptation of test and trace strategies [13].

Since the onset of the SARS-CoV-2 pandemic, multiple new variants of concern (VOC) have emerged as follows [14, 15, 16, 17, 18] :

- Alpha (B.1.1.7): It was first detected in the United Kingdom (UK) in late December 2020. This variant contains several key mutations in the spike protein that distinguish it from the original Wuhan strain. The WHO monitored the spread of Alpha variants containing an additional E484K mutation, which may help the virus to escape the body’s immune defenses by evading neutralizing antibodies generated through vaccination or previous infection.
- Beta (B.1.351): This variant was first reported in South Africa in December 2020. It was more transmissible than previous variants and was considered a concerning variant in terms of reducing neutralization by antibodies generated through previous infection, as well as vaccine efficacy. It implies that people who have already recovered from COVID-19 are at risk of being reinfected, or vaccination may be less effective against it.
- Gamma (P.1): It was detected in Brazil in early January 2021. This variant is associated with increased transmissibility due to its ability to evade humoral immunity and cause reinfections. It was estimated to result in virus levels 3 − 4 times higher than earlier variants and responsible for 1.1−1.8 times more deaths.
- Delta (B.1.617.2): This variant was first reported in India in December 2020. It is estimated to be more contagious than the Alpha variant, and roughly twice as transmissible as the original Wuhan strain of SARS-CoV-2. Epidemiological studies have highlighted that mutations in the spike of the Delta variant may increase infectivity and reduce neutralization to sera from individuals infected with prior variants. These escape mutations are implicated in reinfection, but the observed reduction in the effectiveness of immunization has been modest, with continued strong protection against hospitalization, severe disease, and death.
- Omicron (B.1.1.529): It was initially detected in South Africa in November 2021. It has a large number of mutations, including in the receptor-binding domain of the spike protein which causes a 13-fold increase in viral infectivity and is 2.8 times more infectious than the Delta variant.

As these variants circulated around the globe, a central topic of discussion emerged about the future directions of the evolution of the virus [19]. A motivation of our work is the paper by G. Lobinska et al. [20], where the authors studied the evolution of resistance to COVID-19 vaccination in presence of social distancing. They have derived a formula for the probability of the emergence of vaccine resistance over time for a model with two strains, WT (wild-type virus) and vaccine-resistant mutant virus (MT). In their simulations the social activity level (contact numbers) is adjusted such that the number of infected individuals remains constant in time (i.e. the effective reproduction number is modulated to one). They found that under slow vaccination, resistance is more likely to emerge even if social distancing is maintained, while in the case of rapid vaccination, the emergence of mutants can be prevented if social distancing is observed during vaccination.

Here we construct a somewhat similar model, but one that includes the emergence of *n* (*n* ∈ ℕ) new strains via mutations. Our focus is to investigate the emerging patterns with specific cross-immunities between the strains when disease spread is constrained by dynamically changing social distancing.

This work is outlined as follows: Section 2 introduces our general model. Then two different scenarios are proposed for this model. The one-way cross-immunity scenario is considered in Section 3. In this scenario, recovered individuals gain immunity against the strain that infected them and all older strains, but they are still susceptible to newly emerging strains. We compute the time-varying effective reproduction number for each strain (Subsection 3.3) and discuss the patterns of a consecutive wave-obtained numerical simulations. In Section 4 the second scenario is introduced and compared with the first one. Here recovered individuals gain immunity only against the one strain that infected them, and they are assumed to be susceptible to both new and previous strains. This situation is leading to more complex dynamics compared to the first scenario. Finally, we compare our results with the collected data on COVID-19 variants in the Netherlands in Section 5.

## 2 Model description

We construct a compartmental model to describe a general model of an infectious disease with multiple variants. In our model, the population *N* is divided into the following three main classes, tracking the disease status of individuals: *S* denotes susceptible individuals, i.e. those who can be infected by the disease, *I* denotes infected individuals, and *R* represents the population of recovered individuals. This classical *SIR*-setting is extended now by multiple strains. We use the notation *I*_*k*_ for the class of individuals who are infected by strain *k* (*k* = 1, 2, …, *n*). Upon recovery, individuals move to compartment *R*_*k*_. As a simplification, for those who went through more than one infection, possibly by different strains, their immune status is associated with the last recovery. Hence, individuals in *R*_*k*_ are those who recovered from infection *k*, and they all have the same type of immunity, irrespective of whether they might have been infected in the past by some other strains. The model neglects any changes in the population due to birth, death, or migration during the period of consideration. Thus, the total population (*N*) would be invariant, and for all *t*,

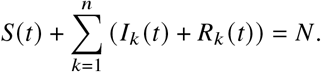

We introduce the general cross-immunity matrix *C* = *c*_*i, j*_ (*i, j* = 1, 2, …, *n*) as the relative immunity to strain *i* for an individual that has been last recovered from strain *j* and thus being in *R*_*j*_ . With this notation, *c*_*i, j*_ = 0 denotes full immunity of *R*_*j*_ individuals to strain *i*, and *c*_*i, j*_ = 1 means no immunity at all (full susceptibility) to the strain. If the value of *c*_*i, j*_ is between 0 and 1, then there is partial immunity to strain *i*. In the model equations, the coefficients *c*_*i, j*_ will appear as a reduction factor in the transmission rate between compartments *R*_*j*_ and *I*_*i*_. The strains are assumed to share the same epidemiological parameters (transmission and recovery rates), they only differ in terms of population immunity against them.

The proposed model is governed by the following system of differential equations:

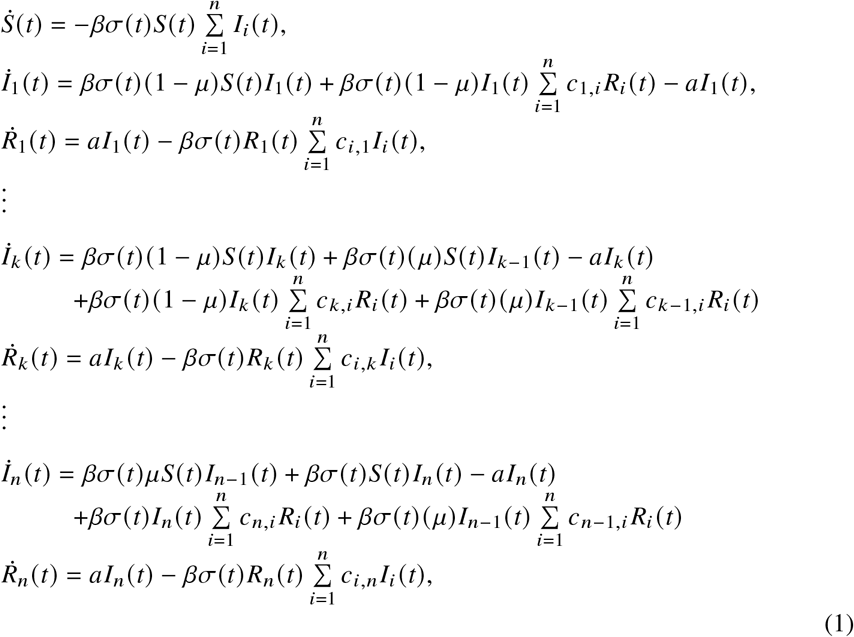

where *k* = 2, 3, …, *n* − 1.

In the above system, parameters *β* and *a* are the average transmission rate and recovery rates, respectively. In this model, a mutation occurs (with a small mutation probability *μ*) when exposure to an infected individual with strain *i* results in an infected individual transmitting further strain *i* + 1. Hence, the mutation is assumed to be sequential, always producing the next variant, where *i* = 1, 2, …, *n*. Moreover, recovered individuals from strain *i* are fully protected from *I*_*i*_ (those infected with strain *i*), and susceptible or immune to others as given by the elements of matrix *C*.

In System (1), parameter *σ* = *σ*(*t*) is a social distancing measure that varies over time and ranges in [0, 1], representing the reduction in contacts between individuals compare to baseline. Here, *σ* = 1 indicates no social distancing, and *σ* = 0 means complete lockdown. Social distancing has a key role in our model since it is a control measure to keep the infected population within an acceptable level. Denoting by *L* this daily new allowed infection threshold, *σ*(*t*) is adjusted such that the infectious population does not exceed *L*/*a*. (see table 1 for parameter references)

**Table 1.** Parameters and values applied in the simulations. The parameters are set to give *R*_0_ = 3.

| Parameter | Interpretation | Values |
| --- | --- | --- |
| $\beta$ | Transmission rate | $7.5 \times 10^{-7}$ |
| $a$ | Recovery rate ($\text{day}^{-1}$) | 0.25 |
| $\mu$ | Mutation probability | $10^{-6}$ |
| $\sigma$ | Social distancing parameter (reduction in contacts) | $[0, 1]$ |
| $N$ | Total population | $10^6$ |
| $L$ | Number of daily new infected (allowed incidence) | 900, 1500, 3500 |

In the following, we focus on two scenarios for the matrix *C*: *i*) if recovery from strain *i* provides immunity against old strains, then *C* is a triangular matrix (discussed in Section 3), and *ii*) if recovery from strain *i* provides immunity only against *i*, but such individuals remain susceptible to both earlier and later strains, then *C* is a diagonal matrix (discussed in Section 4).

## 3 Scenario One: One-Way Cross-Immunity Towards Earlier Variants

### 3.1 Model equations and the cross-immunity matrix

In this case, recovered individuals from strain *i* (*i* = 1, 2, …, *n*) are fully immune to any strain *j*, where *j* ≤ *i*, but have no immunity to the upcoming strains *j*, where *j* > *i*. The cross-immunity matrix below, *C*, represents the above description. For instance, it could be understood that the recovered individual from strain three is immune to strain three and the previous strains as well, (*c*_3, *j*_ = 0, *j* = 1, 2, 3), but they are still susceptible to the new strains, therefore can be infected by them at the same rate as susceptible individuals (*c*_3, *j*_ = 1, *j* = 4, …, *n*). Hence,

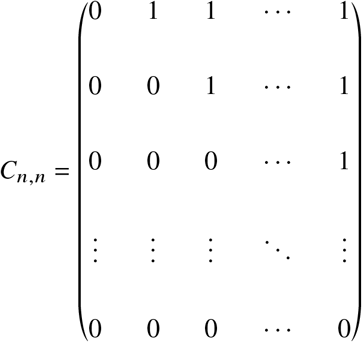

is an upper triangular cross-immunity matrix, with zeros in the diagonal as well. In order to derive the model equations, keep in mind a series of assumptions:

- each strain can infect susceptible individuals *S*;
- recovered individuals from strain *i* are susceptible to strain *j*, *j* > *i*, and immune to previous ones;
- individuals infected by strain *i* can transmit strain *i* + 1 due to mutation with mutation probability *μ*.

Based on these assumptions, the system of differential equations for the model is given by:

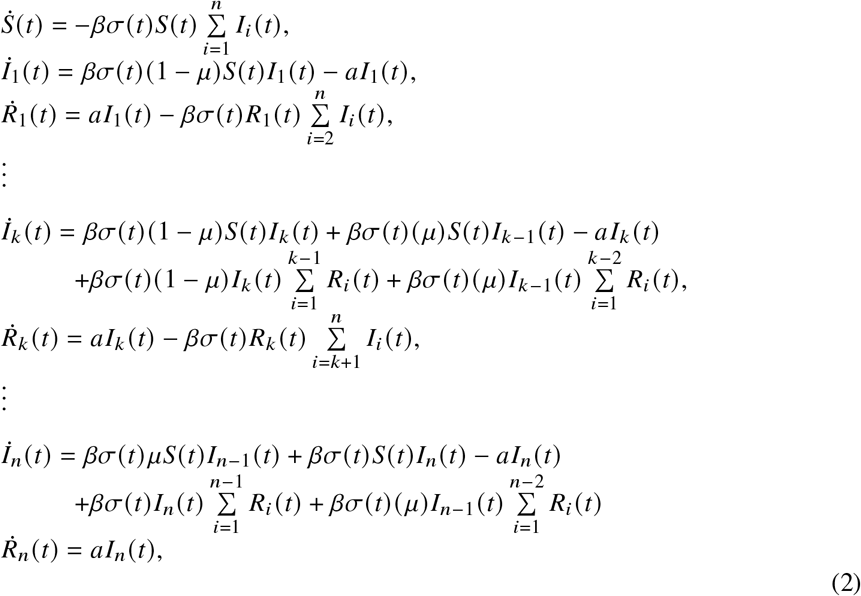

where *k* = 2, 3, …, *n* − 1.

### 3.2 Epidemiological dynamics

We have numerically solved System (2), and Figure 1 shows the number of infected individuals in time for each strain. Initially, we consider 1000 infected individuals by strain 1, while the whole population is considered *N* = 10^6^. In the early stage of the epidemic, there is no social distancing, therefore, *σ* = 1. Then, when the infected population reaches the level *L*/*a* (corresponding to daily incidence *L*), we apply social distancing to the population and set it so that the infected population remains this fixed number, 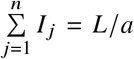. Under this assumption, the effective reproduction number during these times would be equal to one (it will be explained in detail in the next subsection). The number of infections of the first strain decreases due to recovery, and since we considered a fixed infected population, it can be observed in Figure 1 that the newly emerged strains will sequentially replace the previous ones in dominance, having a similar slope to the old declining strains.

**Fig. 1.**
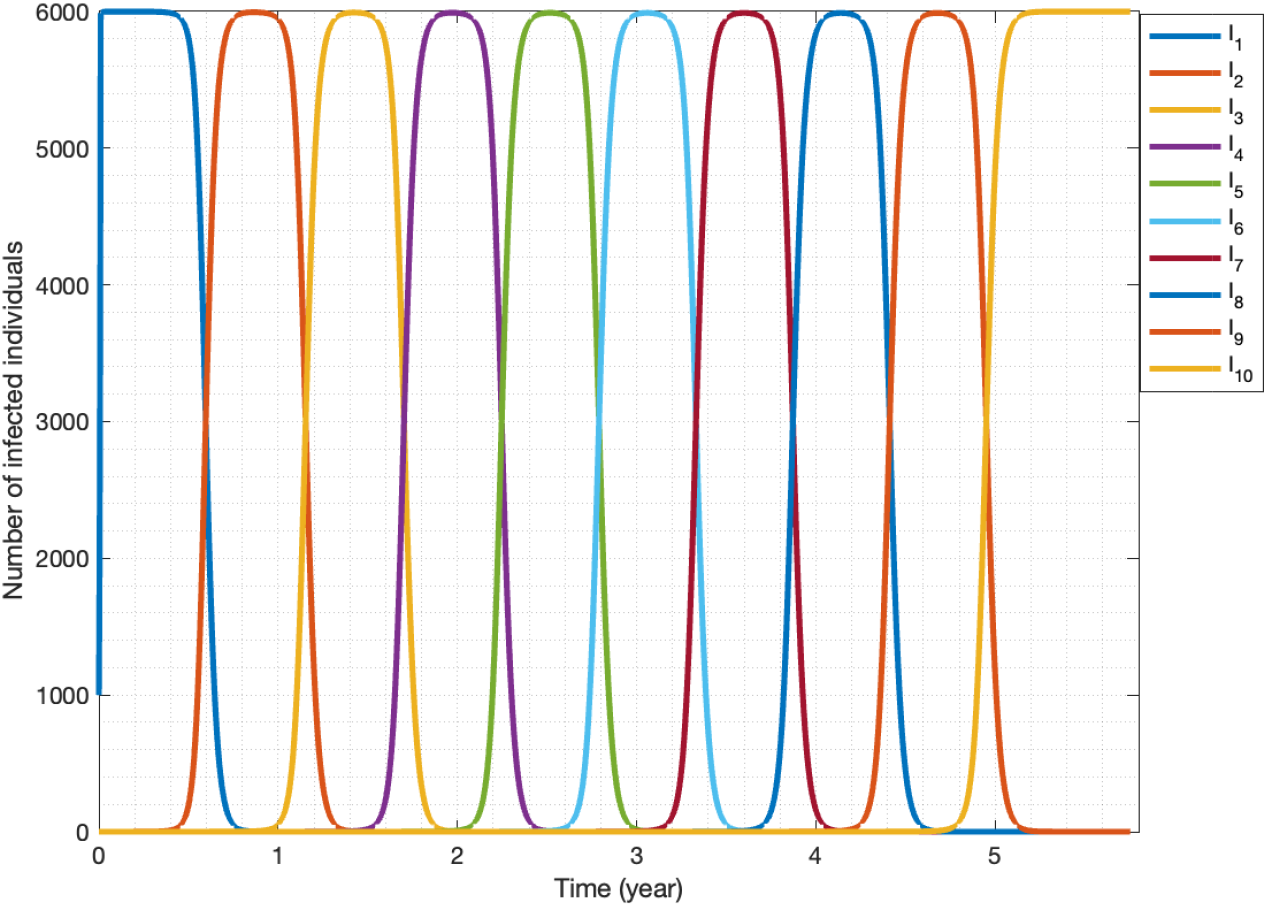
Infections behavior over time of scenario 1; *N* = 10^6^, *L* = 1500, *a* = 0.25, *β* = 7.5 × 10^−7^.

Figure 2 represents how the number of allowed new infections per day affects the imposed social distancing over time: a high number of newly infected individuals per day leads to higher *σ*(*t*), corresponding to milder interventions for social distancing. It is also noticeable that when a dominant strain is in decline, we can relax the measures to some extent, but later we need to make it more stringent again, giving rise to an oscillatory pattern as newer and newer strains are taking over.

**Fig. 2.**
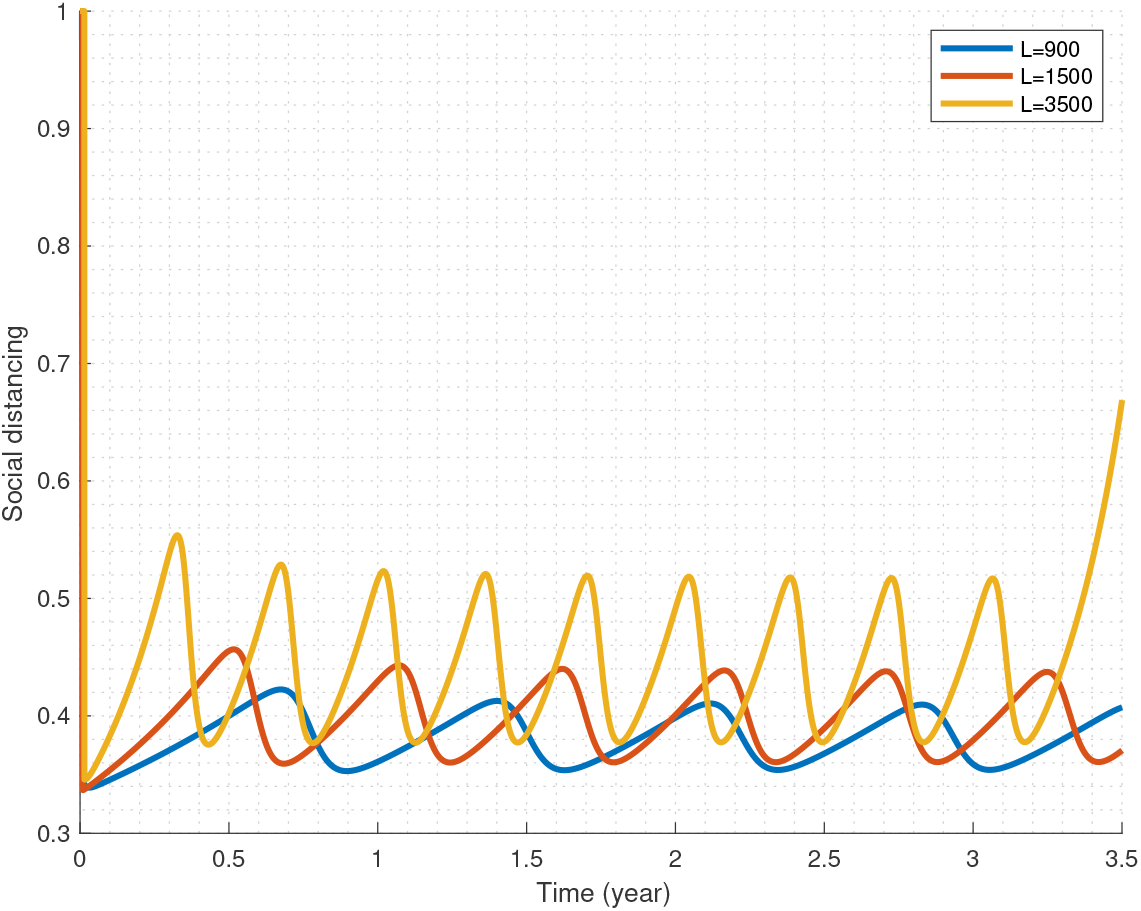
Relation between social distancing parameter, *σ*(t), and *L* for ten strains of first Model.

Looking at Figure 1, a natural question is whether the dominance period of newer strains is the same as earlier ones since on the graph they look very similar. Hence, we compare the dominance periods in the population for each strain with three different values of *L*. The interesting outcome is represented in Figure 3. This figure reflects the fact, on one hand, that the more new individuals are allowed to be infected by a strain, the shorter that strain remains in dominance and the faster it fades out. To calculate the period of dominance of a variant in the population, the distance between two different times is calculated when the number of infected individuals with each strain is *L*/(2*a*) . As seen in Figure 1, the number of people infected with strain *i* (*i* = 1, 2, …, *n*) reaches *L*/(2*a*) at two different times, when strain *i* is emerging (strain *i* − 1 is dying out) and fading out (strain *i* + 1 is emerging). Then, the dominance period is defined as the time difference between these two points, i.e. the time duration for a strain having a higher number of infections than *L*/(2*a*) . As Figure 3 shows, in this scenario, each subsequent strain has a shorter period of dominance than the previous ones. On the other hand, higher allowed incidence corresponds to shorter dominance periods and faster emergence of novel strains.

**Fig. 3.**
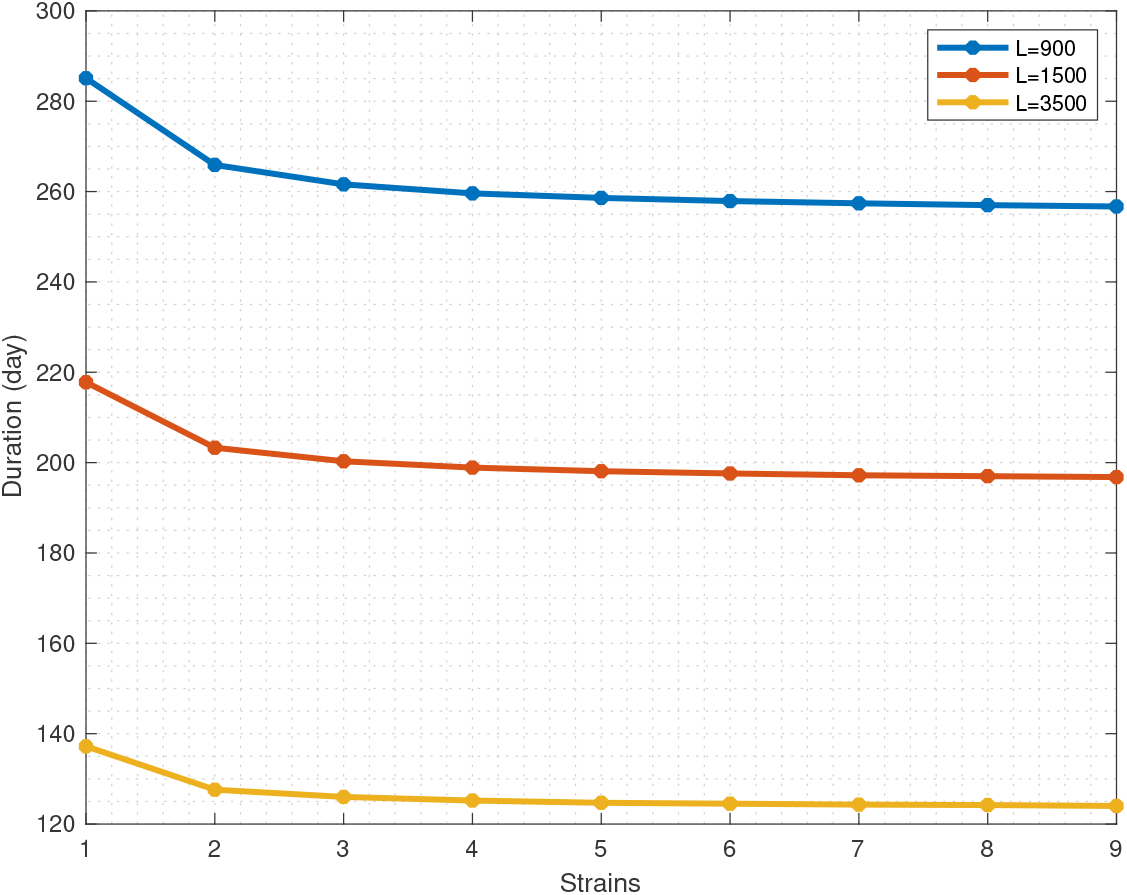
Relation between new infections per day (*L* (*t*)) and duration of persistence in the population.

### 3.3 Reproduction Numbers

Figure 4 exhibits the effective reproduction number for each strain as it changes over time. Initially, all strains can potentially infect the entire population, and as time goes by each strain infects fewer individuals due to recovery from the infection by that strain. New mutations can still potentially infect every individual in the population since recovered individuals from old strains are susceptible to new ones. Since, according to our basic assumption, the social distance parameter is manipulated so that we have a fixed number of infected populations 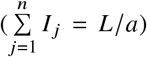, the effective reproduction remains at 1 for model 1 (dashed line in Figure 4).

**Fig. 4.**
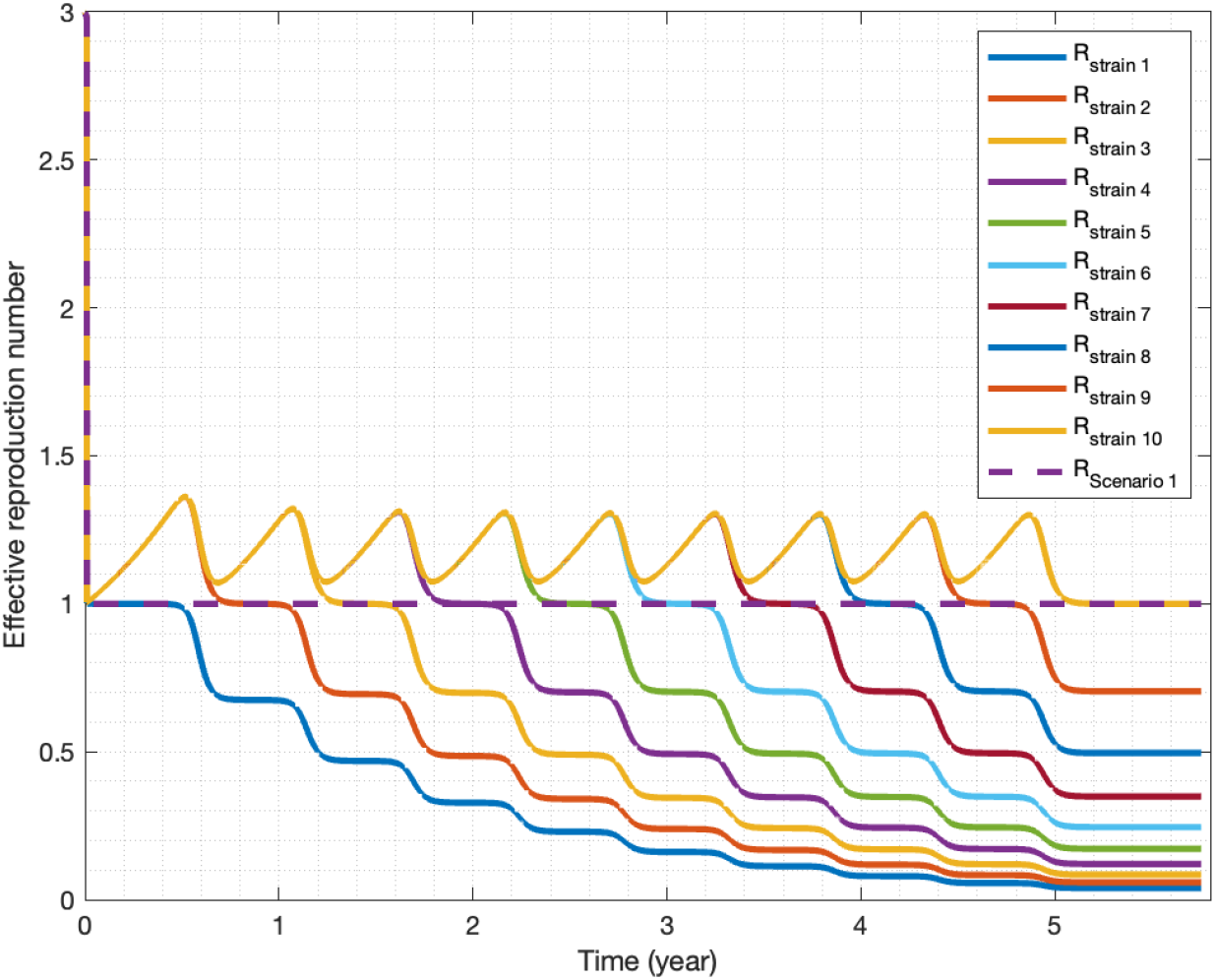
Effective reproduction number for each strain of system 1.

The overall effective reproduction number is the weighted average of all strain reproduction numbers, weighted by the number of infected individuals by that strain. To calculate these specific reproduction numbers, we create the next generation matrix *FV*^−1^, where matrices *F* and *V* are defined as follows [21]:

Let *I* = (*I*_1_, *I*_2_, …, *I*_*n*_)^*T*^ is the number of individuals in infection compartments in System (2). We rewrite the corresponding equations in the form of *İ*_*i*_ = ℱ_*i*_ (*I*) −𝒱_*i*_ (*I*) for *i* = 1, 2, …, *n*. Here, ℱ_*i*_ is the term for the appearance of new infections in compartment *i*, and 𝒱_*i*_ is the rate of transitions between compartment *i* and other infected compartments. Define non-negative matrix 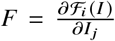 and non-singular matrix 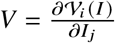 for 1 ≤ *i, j* ≤ *n*. This way, we obtain

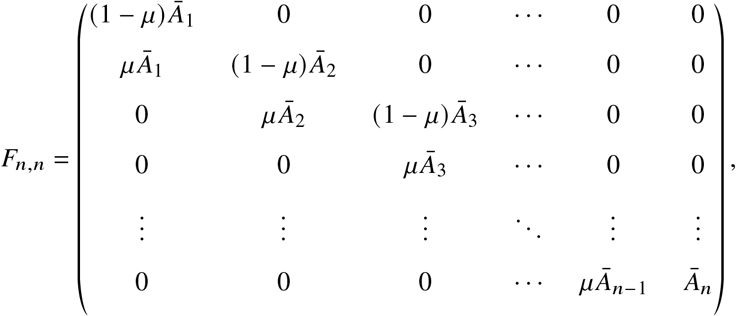

and

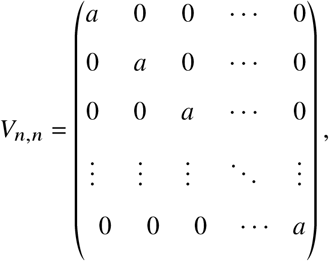

where Ā_1_ = *βσS* and 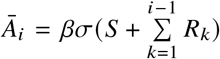 for *i* = 2, 3, …, *n*. At the next step, we generate the next generation matrix *FV*^−1^ as

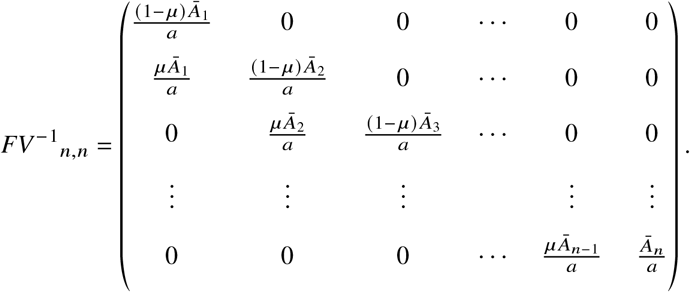

The eigenvalues of the next generation matrix give us the effective reproduction numbers for each strain, varying with time. The eigenvalues

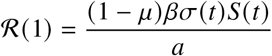

and

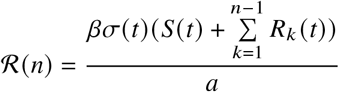

are effective reproduction numbers corresponding to the first and last strains, respectively, and

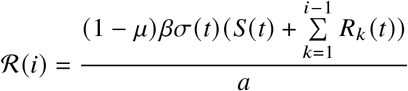

is the effective reproduction number for infection compartments *I*_*i*_, *i* = 2, 3, …, *n* − 1. Moreover, the overall effective reproduction number for the first scenario of our model can be obtained by

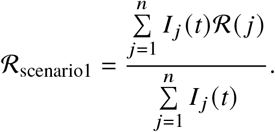

Note that in general, the next generation matrix is calculated at a steady state. Here, to calculate the effective reproduction numbers to every *t*, for a given *t* we freeze the values of *S*(*t*), *σ*(*t*), *R*_*k*_(*t*) for the time period the subsequent infected generation being created, ignoring short term changes in these values and treating them as steady states, and we perform the same formal calculations as in the classical case in a true steady state.

## 4 Second Scenario: Absence of Cross-Immunity

### 4.1 Model equations and the cross-immunity matrix

Now, let us assume protection upon recovery from one strain holds only against that particular strain, thus they are susceptible to old and new strains, e.g. in the cross-immunity matrix below the diagonal elements are zero which represents full immunity to the strain that recovered individuals have been recovered from, and other elements are one which conveys no protection against the novel and earlier strains. Then,

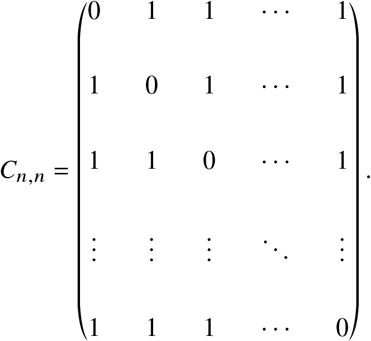

The second scenario is based on the assumptions that are listed below:

- each strain can infect susceptible individuals *S*;
- recovered individuals from strain *i* are only resistant to this strain and not protected against other strains;
- individuals infected by strain *i* can transmit strain *i* + 1 due to mutation.

The dynamics of the epidemic model we described can be summarized by the following equations for *k* = 2, 3, …, *n* − 1:

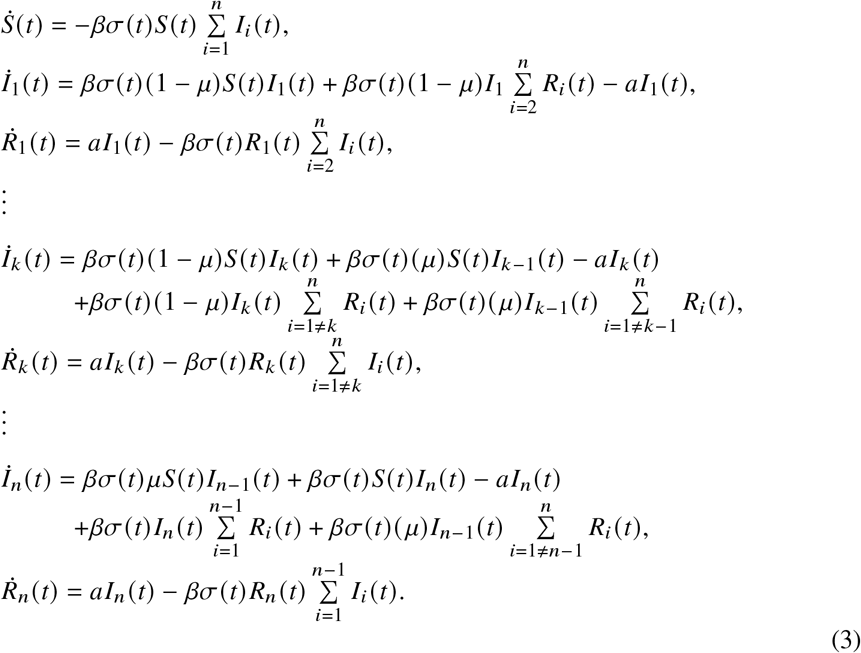

### 4.2 Epidemiological dynamics

In this case, unlike in Scenario 1, recovered individuals are not immune to old variants, so the infected populations in Figure 5 do not have as regular behavior as those in Figure 1. As we can see in Figure 5, for each strain the infections settle around the value 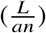 over time, In other words, for large *t*:

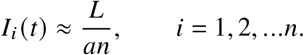

**Fig. 5.**
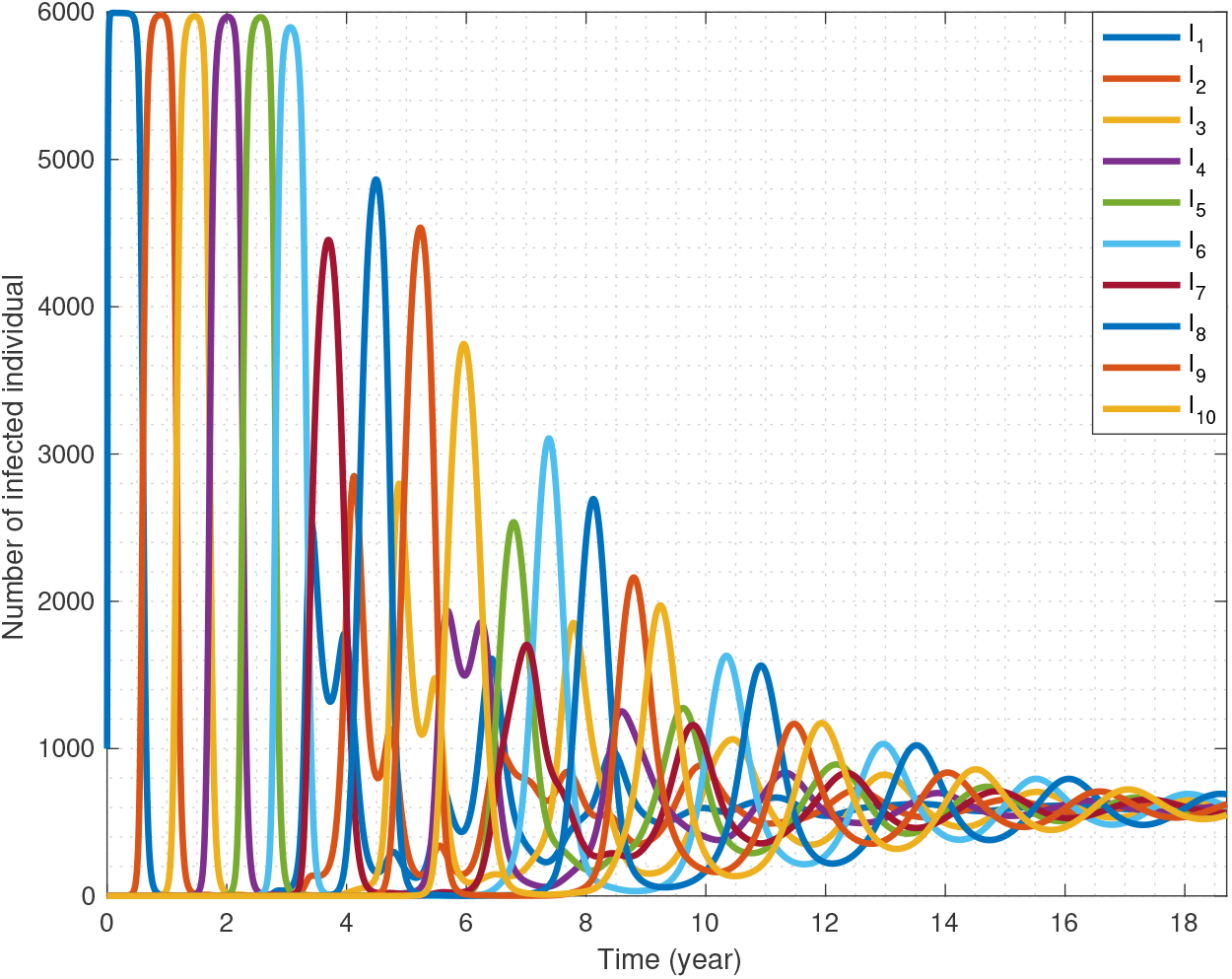
Infected population of each strain of Scenario 2, that are approaching *L*/(*an*).

This can be explained intuitively as follows: In the first scenario, we have susceptibility only to the new strains, that is, recovered people cannot be infected with old strains and there is no mutation from new strains to the previous strains, so the earlier strains can not compete with newer ones and converge to zero, while the new ones rise to *L*/*a*. However, in Scenario 2, each strain infects all recovered individuals, who were recovered from new and old strains (except from the very same strain), hence their potential pools are equalized. Moreover, the total amount of infected is forced to be *L*/*a*, which is now distributed among *n* equally competitive strains.

Consequently, the effective reproduction numbers go rounding around one since the number of infected individuals neither increases nor decreases during these times (Figure 6).

**Fig. 6.**
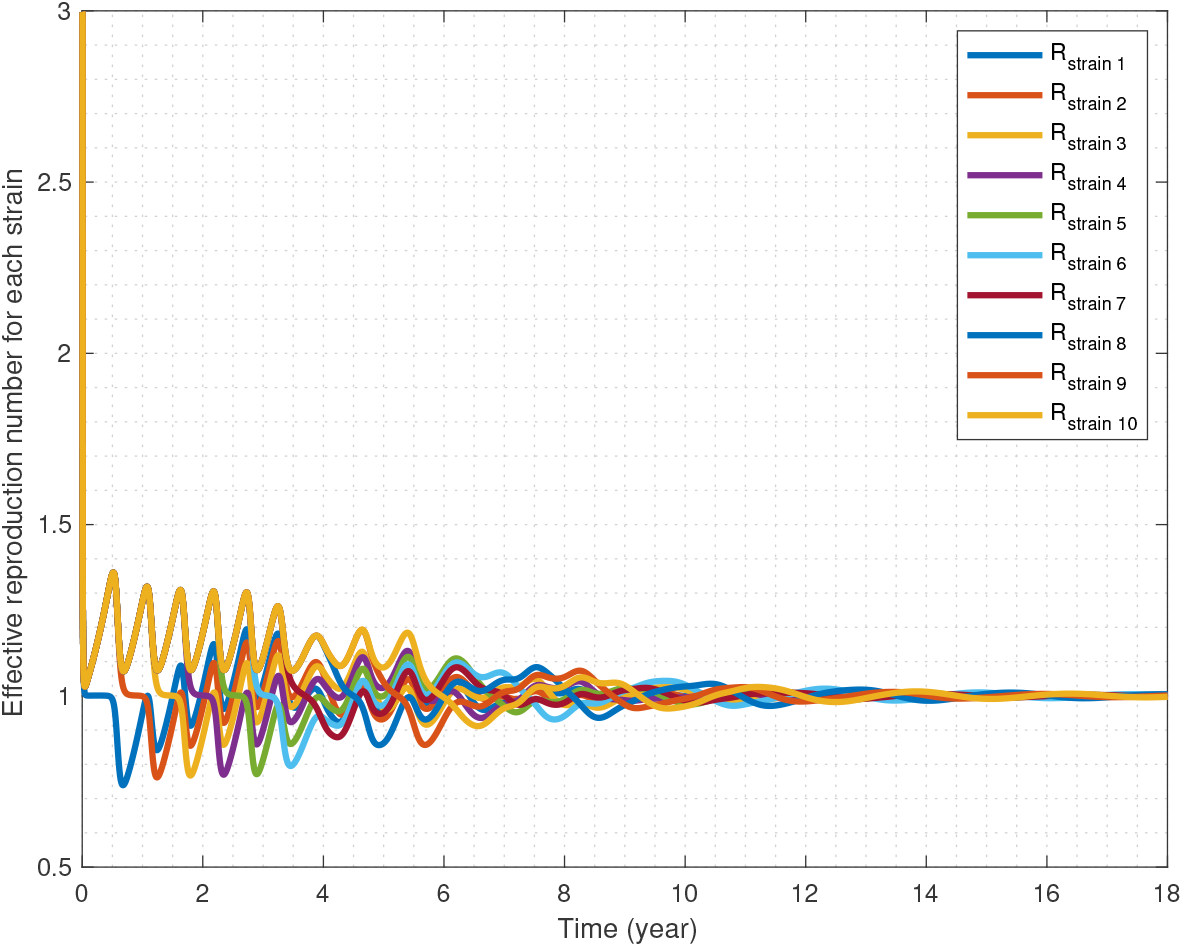
Effective reproduction numbers for each strain approach one as the infected population approaches *L*/(*an*).

### 4.3 Effective Reproduction Numbers

To compute the effective reproduction number for each strain in this scenario, we proceed as in Subsection 3.3: we first generate the next generation matrix, and then calculate its eigenvalues which are the effective reproduction numbers corresponding to each strain.

We rewrite the equations for the infected compartments *I* = (*I*_1_, *I*_2_, …, *I*_*n*_)^*T*^ in System (3), as

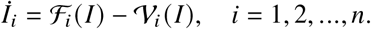

ℱ_*i*_ and 𝒱_*i*_ are defined the same way as in Subsection 3.3. Define the non-negative Matrix 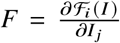, and the non-singular matrix 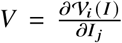 for 1 ≤ *i, j* ≤ *n* as follows:

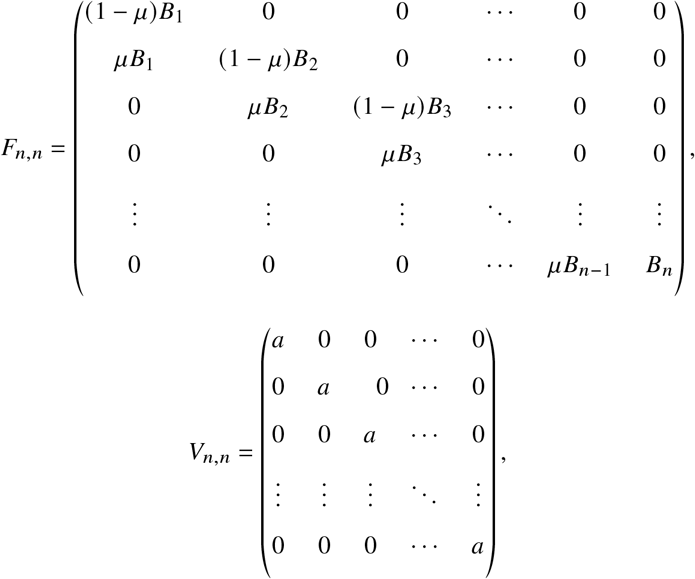

Where 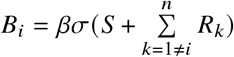 for *i* = 1, 2, …,*n*. Then we create the matrix *FV*^−1^

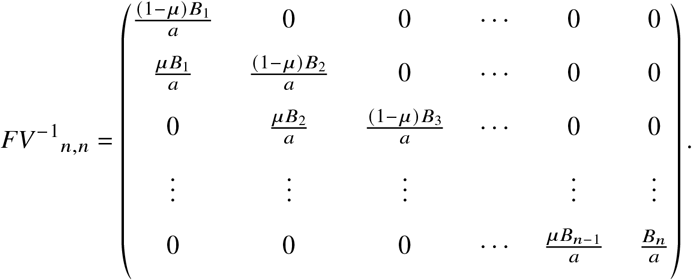

The eigenvalues of this matrix give us the effective reproduction numbers for each strain. In particular, the eigenvalue

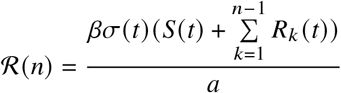

is the effective reproduction number that corresponds to the last strain, and

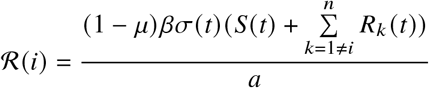

is the effective reproduction number corresponding to strain *i, i* = 1, 2, …, *n* − 1.

## 5 Discussion

The way in which novel mutations of infectious diseases, particularly COVID-19, emerge and persist in the population prompts us to implement a general model with multiple strains emerging via mutations. For the sake of simplicity, here we assumed that the strains have identical epidemiological parameters, and they differ only in the target population which they can infect, and this is determined by a cross-immunity matrix. For the purposes of this paper, we considered two different scenarios which focus only on immune evasion of newer strains emerging by virus mutations. Recovery from a strain in the first scenario confers full protection against previous strains and no immunity to the novel strains. From the simulation results, we can observe a highly structured sequential behavior of strain replacement where newer strains predominate over older strains, but last in the population for an ever shorter period of time. However, since in the second scenario, immunity upon recovery holds only against a given strain and none of the others, the population infected by each strain has an erratic pattern over time, with recurrence of past strains, and co-circulation of many strains without any of them being clearly dominant. In the following example, we compare our results with the reported data on the coronavirus SARS-CoV-2 variants in the Netherlands.

Figure 7 represents the frequency of COVID-19 variants since 2021 in the Netherlands, based on the data from [22]. In this graph, variants Alpha and Delta are emerging after each other and are fully dominant, showing a very similar picture to our Scenario 1, where even the periods of dominance are of similar length in the model and in the data. But almost after June 2022, most of the variants of SARS-CoV-2 are from the Omicron lineage, from sub-variants BA.1 through BQ.1. While the BA.5 sub-variant is still dominant, some new sub-variants such as BQ.1 are gaining. The recently emerging Omicron sub-variants are alarmingly immune evasive [23], and they could further compromise the effectiveness of current COVID-19 vaccines, as well as curtailing prior natural immunity. This, in turn, could increase cases of infections and reinfections. The recent situation, regarding the cross-immunity between novel strains is more similar to our Scenario 2, where, after a while we can see irregular circulation patterns in Figures 5, just as in 7. Thus, this phenomenon is more similar to the Omicron-era Netherlands data. This confirms that our model captures some of the essence of the variant dynamics that we have seen during COVID-19.

**Fig. 7.**
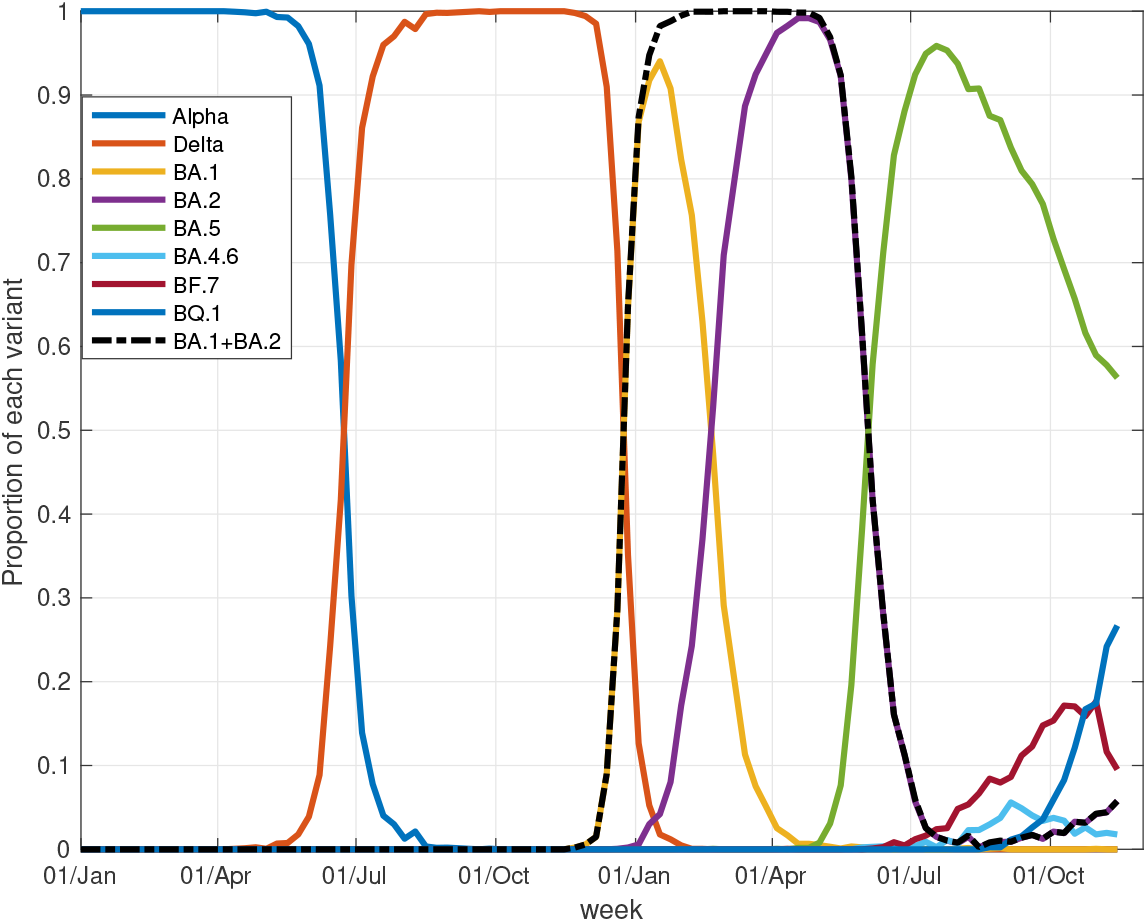
Variants of the coronavirus SARS-COV-2 in the Netherlands from 30/11/2020 to 14/12/2022.

## Acknowledgements

GS has received funding from the European Union’s Horizon 2020 research and innovation programme under grant agreement No 955708, EvoGamesPlus. GR was supported by Hungarian grants NKFIH KKP 129877, RRF-2.3.1-21-2022-00006, and TKP2021-NVA-09.

